# The Ca^2+^-activated cation channel TRPM4 is a positive regulator of pressure overload-induced cardiac hypertrophy

**DOI:** 10.1101/2020.12.21.423727

**Authors:** Yang Guo, Ze-Yan Yu, Jianxin Wu, Hutao Gong, Scott Kesteven, Siiri E. Iismaa, Andrea Y. Chan, Sara Holman, Silvia Pinto, Andy Pironet, Charles D. Cox, Robert M. Graham, Rudi Vennekens, Michael P. Feneley, Boris Martinac

## Abstract

Pathological left ventricular hypertrophy (LVH) is a consequence of pressure overload caused by systemic hypertension or aortic stenosis and is a strong predictor of cardiac failure and mortality. Understanding the molecular pathways in the development of pathological LVH may lead to more effective treatment. Here, we show that the transient receptor potential cation channel subfamily melastatin 4 (TRPM4) ion channel is an important contributor to the mechanosensory transduction of pressure overload that induces LVH. In mice with pressure overload induced by transverse aortic constriction (TAC) for two weeks, cardiomyocyte TRPM4 expression was reduced, as compared to control mice. Cardiomyocyte-specific TRPM4 inactivation reduced by ~50% the degree of TAC-induced LVH, as compared with wild type (WT). In WT mice, TAC activated the CaMKIIδ-HDAC4-MEF2A but not the calcineurin-NFAT-GATA4 pathway. In TRPM4 knock-out mice, activation of the CaMKIIδ-HDAC4-MEF2A pathway by TAC was significantly reduced. However, consistent with a reduction in the known inhibitory effect of CaMKIIδ on calcineurin activity, reduction in the CaMKIIδ-HDAC4-MEF2A pathway was associated with partial activation of the calcineurin-NFAT-GATA4 pathway. These findings indicate that the TRPM4 channel and its cognate signalling pathway are potential novel therapeutic targets for the prevention of pathological pressure overload-induced LVH.

**Significance statement:** Pathological left ventricular hypertrophy (LVH) occurs in response to pressure overload and remains the single most important clinical predictor of cardiac mortality. Preventing pressure overload LVH is a major goal of therapeutic intervention. Current treatments aim to remove the stimulus for LVH by lowering elevated blood pressure or replacing a stenotic aortic valve. However, neither of these interventions completely reverses adverse cardiac remodelling. Although numerous molecular signalling steps in the induction of LVH have been identified, the initial step by which mechanical stretch associated with cardiac pressure overload is converted into a chemical signal that initiates hypertrophic signalling, remains unresolved. Here, we demonstrate that the TRPM4 channel is a component of the mechanosensory transduction pathway that ultimately leads to LVH.

## Introduction

Pathological left ventricular hypertrophy (LVH) is the most powerful independent predictor for cardiovascular mortality (1, 2). It occurs in response to two very common clinical conditions: systemic hypertension and aortic valve stenosis. It manifests as increased cardiomyocyte volume and weight (3–6), which results in increased heart mass, particularly left ventricular (LV) mass. Although pathological LVH commonly occurs as a response to increased cardiac wall stress, sometimes termed “compensatory hypertrophy”, it is now well established that the effects of pathological LVH are deleterious for heart function, leading to increased cardiac failure and death (1, 2). So far, the only treatment for this condition is lowering elevated blood pressure or replacing a stenotic aortic valve. However, these treatments cannot completely reverse the pathological effects on the myocardium once LVH is established. Consequently, understanding the molecular mechanisms underlying pathological LVH may lead to therapies directed at preventing, inhibiting, or reversing pathological LVH, and reducing its associated morbidity and mortality.

The development of pathological LVH depends on upstream stimuli, such as mechanical forces (e.g., pressure overload) or neuroendocrine hormones (e.g. angiotensin II), and distinct downstream signalling mechanisms (7–11). Importantly, a large body of work implicates intracellular Ca^2+^ levels and subsequent activation of Ca^2+^-calmodulin dependent signalling pathways, such as the calcineurin-NFAT-GATA4 axis, in the induction of pathological LVH (12, 13) (14–19). Gq-coupled receptors are thought to play an important role in the induction of pathological LVH in response to both neurohumoral stimulation (11, 20, 21) and mechanical forces, such as the increase in LV afterload induced by experimental aortic constriction (22). Once activated, Gq-coupled receptors are thought to then activate the calcineurin-NFAT-GATA4 pathway (14–19).

Our previous experimental work, however, has demonstrated that although Gq-coupled receptors and the calcineurin-NFAT-GATA4 pathway are essential for the induction of LVH in response to angiotensin II, neither are required for the induction of LVH in response to transverse aortic arch constriction (TAC), the most common experimental model of LV pressure overload (23), and one not associated with activation of the renin-angiotensin system (24). In contrast to the lack of activation of the calcineurin-NFAT-GATA4 pathway with TAC, an alternative Ca^2+^-calmodulin dependent signalling pathway, the CaMKII-HDAC-MEF2 pathway (25–27), is activated in response to TAC (23).

Left unexplained by our previous work, however, is the mechanism by which the CaMKII-HDAC-MEF2 pathway is activated by TAC, given that this activation is not dependent on Gq-coupled receptors. Prime candidates for mediating this mechanism are mechanosensitive ion channels. In cardiac mechano-transduction, where mechanical stimuli are converted into electrical or chemical signals (28, 29), Ca^2+^-dependent ion channels, such as transient receptor potential (TRP) channels (16), act as important modulators of intracellular Ca^2+^ homeostasis (30) and are thought to be unique biosensors that activate specific pathological LVH signalling pathways (31, 32). As a Ca^2+^- and voltage-activated non-selective monovalent cation channel, transient receptor potential cation channel subfamily melastatin 4 (TRPM4) may contribute to an increase in intracellular Ca^2+^ concentration by causing membrane depolarization (33), although we and others (40, 41, 42) have demonstrated that mammalian TRP channels, including TRPM4, are not directly stretch-activated. Consequently, if TRPM4 plays a role in TAC-induced LVH, it acts as an amplifier of the primary Ca^2+^ or voltage signal from a yet to be determined mechanosensitive ion channel, or channels. TRPM4 has been functionally characterized in atrial and ventricular cardiomyocytes, both human and rodent (34–36). Other studies indicate that TRPM4 contributes to both cardiac function and disease development, including cardiac hypertrophy and heart failure (37–41). A previous study using *Trpm4* cardiomyocyte-specific knock-out (*Trpm4*^cKO^) mice has shown that TRPM4 is a negative regulator of angiotensin II-induced cardiac hypertrophy in mice, which involves the calcineurin-NFAT pathway (42). However, whether TRPM4 plays a role in mechanical pressure overload-induced LVH has yet to be determined.

Here, we investigated the role of TRPM4 in pressure overload LVH induced by TAC in homozygous cardiomyocyte-specific *Trpm4* knock-out (*Trpm4*^cKO^) mice (42) as compared to wild type (WT) control mice. We demonstrate that loss of cardiomyocyte TRPM4 significantly attenuates the development of LVH observed in response to TAC in WT mice. Moreover, this effect is associated with reduced activation of the CaMKII-HDAC4-MEF2A pathway.

## Results

### Development of LVH in response to pressure overload at 14 days after TAC in WT mice

As documented in our previous study (23), TAC induced cardiac hypertrophy as a response to LV pressure overload. As expected, LV systolic pressure increased by ~65 mmHg (*p* < 0.001) 14 days after TAC (**Fig. 1A**), whereas heart rate (HR) (**Fig. 1B**), dP/dt_max_ (**Fig. 1C**) and dP/dt_min_ (**Fig. 1D**) remained unaltered. Consistent with 14 days of TAC resulting in a compensated LVH model, body weight (BW, **Fig. 1F**) and lung weight (LW) (**Supplementary Table 1**) remained unchanged in TAC mice compared to sham-operated mice.

**Fig. 1.**
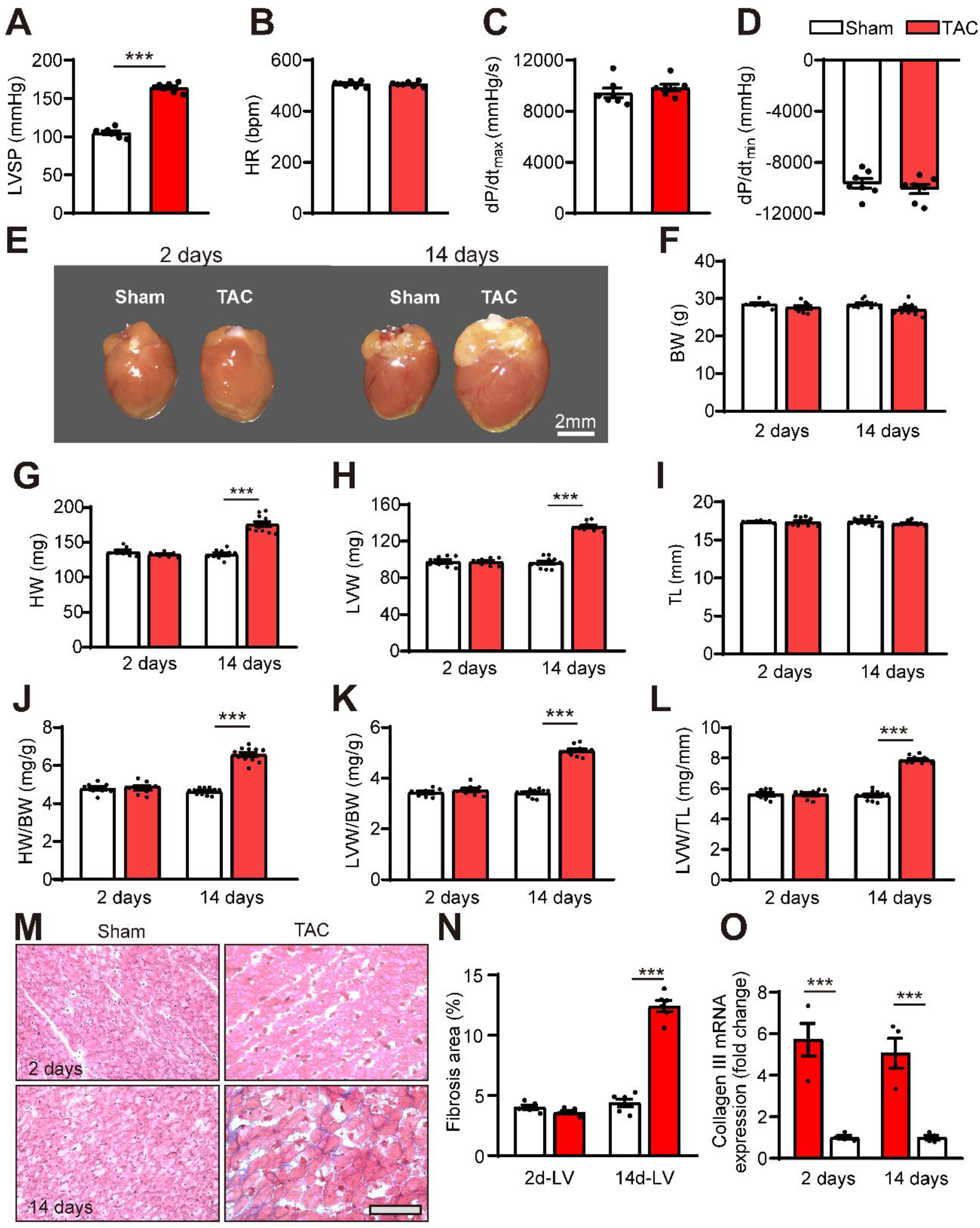
The hypertrophic response to left ventricular pressure overload-induced by TAC. *(A)* LVSP;*(B)* HR; *(C)* dP/dt_max_; *(D)* dP/dt_min_ were measured in wild type (WT) mice 14 days after subjected to TAC versus sham-operated controls (n = 7-11/group). *(E)* Representative photos of hearts from WT mice 2 or 14 days after sham or TAC; *(F)* BW; *(G)* HW; *(H)* LVW; *(I)* TL were measured at the time of sacrifice; *(J-L)* LVH developed 14 days after TAC, as indicated by the ratios of HW/BW, LVW/BW and LVW/TL in WT mice subjected to TAC versus sham-operated controls. *(M and N))* Cardiac fibrosis was evaluated by Masson’s trichrome staining of LV tissue from WT mice subjected to TAC versus sham-operated controls; *(M)* Representative photos; *(N)* Cardiac fibrosis areas were graded (n = 5-6/group). and *(O)* Relative *Col3a1* mRNA expression in WT mice subjected to TAC versus sham-operated controls (n = 4/group). Scale bar = 100 μm. LVSP: Left ventricular systolic pressure; HR: heart rate; dP/dt: first derivative of pressure with respect to time. BW: body weight; HW: heart weight; LVW: left ventricular weight; TL: tibia length; HW/BW: heart weight to body weight ratio; LVW/BW: LV weight to body weight ratio; LVW/TL: LV weight to tibia length ratio. Results are presented as means ± SEM. \*\*\**p* < 0.001, vs. sham-operated groups.

Compared to sham-operated animals (**Fig. 1E-L**), TAC also resulted in significant increases in heart weight (**Fig. 1G**) and size (**Fig 1E**) and in LV mass after 14 but not 2 days (**Fig. 1E, G, H, J-L, K**), and consistent with the development of pathological hypertrophy, TAC was associated with cardiac fibrosis (**Fig. 1M-N**) and enhanced collagen III expression (**Fig. 1O**).

### Early gene markers of pathological hypertrophy-induction in WT mice

Although there was no significant LVH 2 days after TAC (**Fig. 1E-L**), induction of hypertrophy-associated genes [atrial natriuretic peptide (ANP, *Nppa*; 9.4 fold, *p* < 0.001, **Fig. 2A**), brain natriuretic peptide (BNP, *Nppb*; 9 fold *p* < 0.01, **Fig. 2B**) and α-skeletal actin (α-SA, *Acta1*; 4 fold, *p* < 0.01, **Fig. 2C**)] was already evident at this time and expression of these genes remained high at 14 days [ANP, *Nppa* (*p* < 0.001, **Fig. 2D**), BNP, *Nppb* (*p* < 0.001, **Fig. 2E**) and α-SA, *Acta1* (*p* < 0.001, **Fig. 2F**)].

**Fig. 2.**
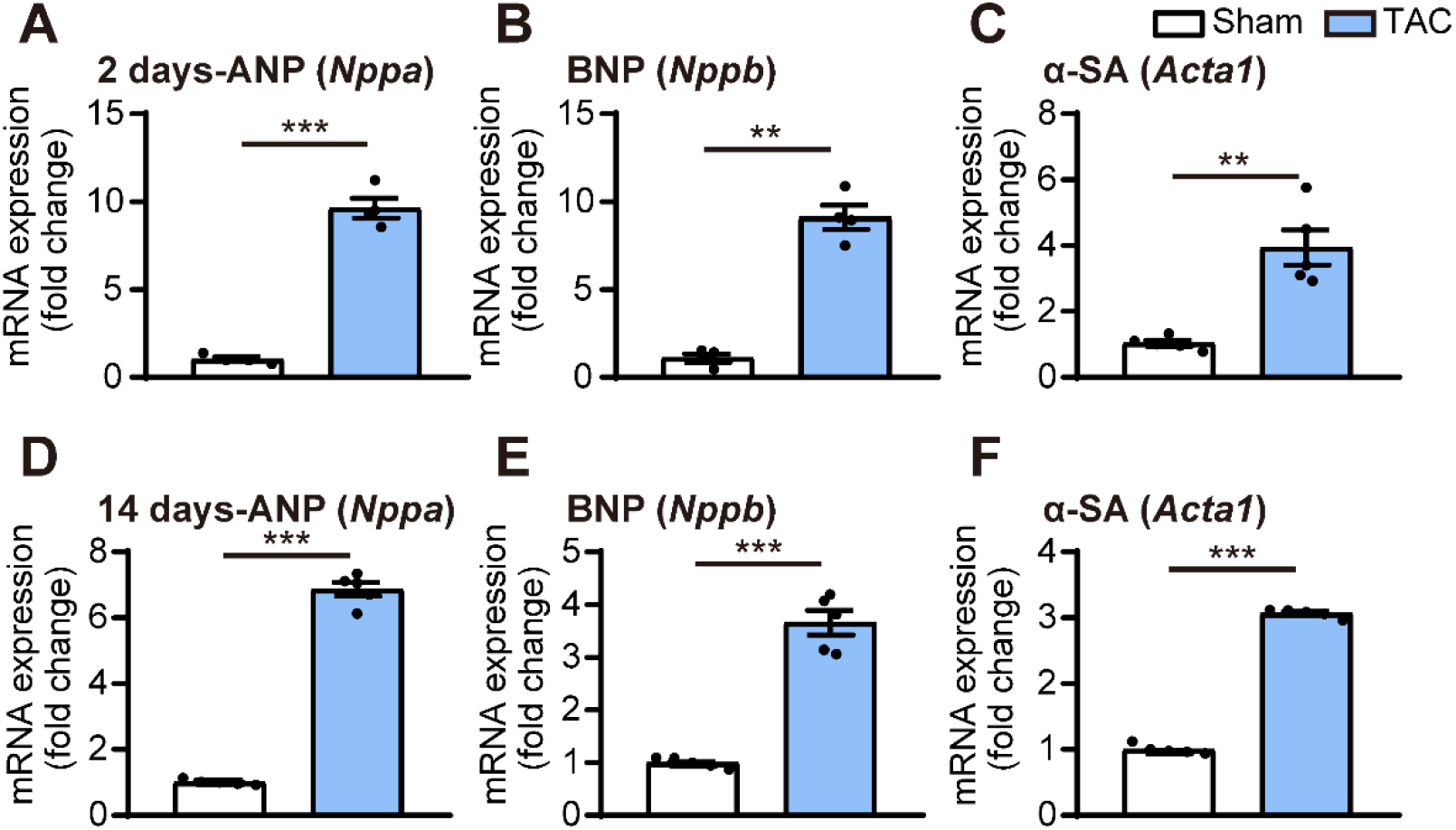
Early markers of LVH induction in response to left ventricular pressure overload. *(A)* Relative mRNA expression of ANP (*Nppa*), *(B)* BNP (*Nppb*) and *(C)* α-SA (*Acta1*) after 2 days of TAC compared to sham (n = 4/group). *(D)* Relative mRNA expression of ANP (*Nppa*), (*E*) BNP (*Nppb*) and *(F)* α-SA (*Acta1*) after 14 days of TAC compared to sham (n = 5/group). The relative mRNA expression was calculated as fold change normalized by GAPDH. Results are presented as means ± SEM. \*\**p* < 0.01, \*\*\**p* < 0.001, compared between sham- and TAC-operated groups.

### TRPM4 expression was downregulated in response to LV pressure overload in WT mice

To examine whether the TRPM4 ion channel is involved in TAC induced LVH, we conducted real-time quantitative PCR (RT-PCR) of LV tissues or isolated LV cardiomyocytes from TAC- or sham-operated hearts. LV tissue and cardiomyocyte *Trpm4* mRNA expression fell by 50 (*p* < 0.001, **Fig. 3A**) and 57% (*p* < 0.001, **Fig. 3B**), respectively, in response to 2 days of TAC and expression continued to be reduced by 30 (*p* < 0.05, **Fig. 3A**) and 40% (*p* < 0.001, **Fig. 3B**), respectively, at 14 days. Consistent with the mRNA changes, LV tissue and isolated cardiomyocyte TRPM4 protein expression also fell significantly, particularly in cardiomyocytes, after 14 days of TAC (Fig. 3C-E).

**Fig. 3.**
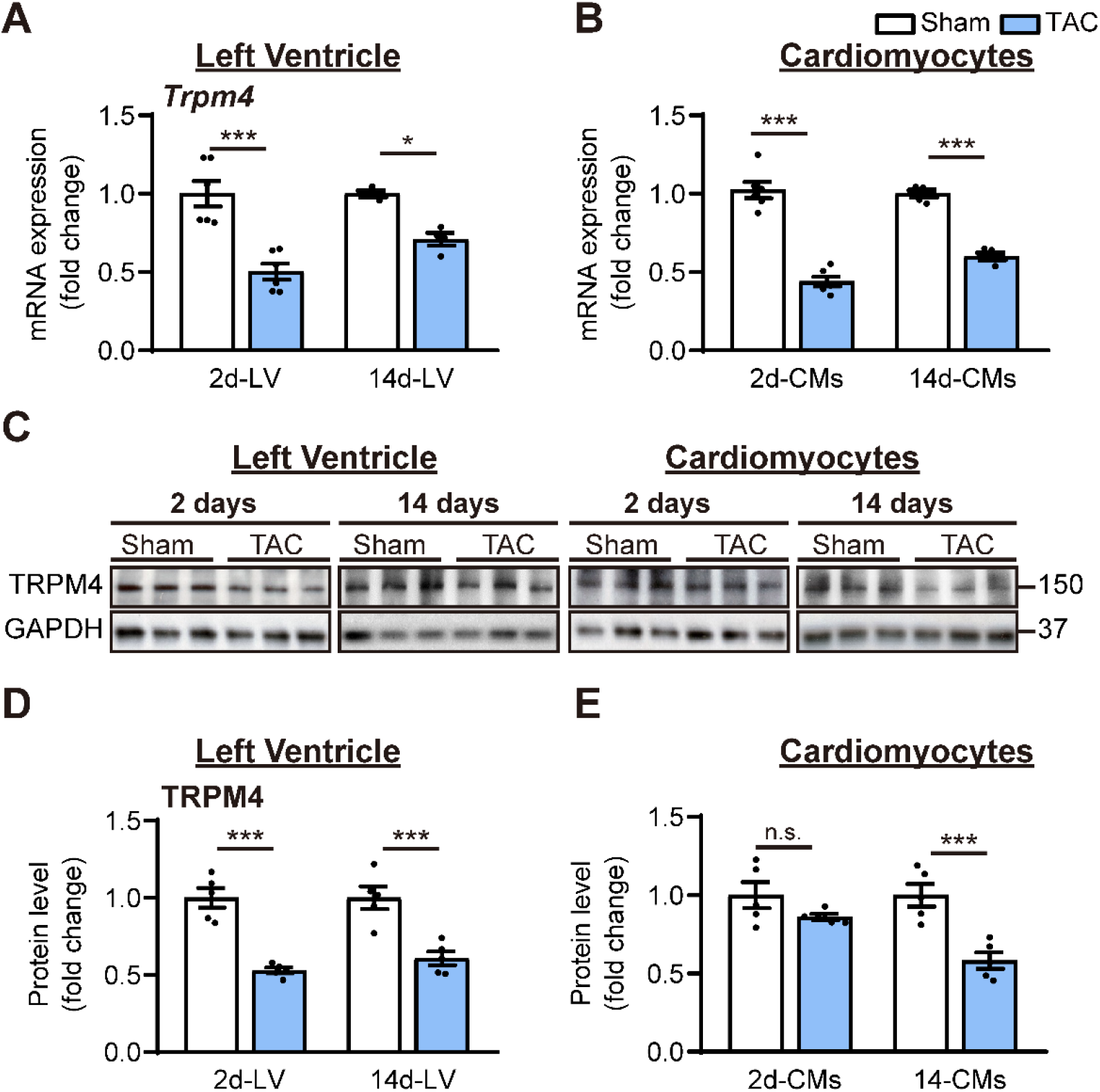
TRPM4 expression was downregulated in response to left ventricular pressure overload. *(A)* Relative mRNA expression of *Trpm4* in LV tissue and *(B)* in left ventricular cardiomyocytes (CMs) after 2 days and 14 days of sham and TAC. *(C)* Representative Western blots of TRPM4 protein expression in LV tissue (*left panel*) and in LV cardiomyocytes (*right panel*). *(D)* Western blots from LV tissues and *(E)* LV cardiomyocytes after 2 days and 14 days of TAC were quantitated for TRPM4 protein expression. Relative TRPM4 mRNA and protein levels in the LV tissue and LV cardiomyocytes were quantified as fold change normalized to GAPDH. Results are presented as means ± SEM. \**p* < 0.05, \*\**p* < 0.01, \*\*\**p* < 0.001 vs. sham-operated groups.

### TRPM4 deficiency decreases the hypertrophic response to TAC-induced pressure overload

To further investigate the role of TRPM4 channels in pressure overload-induced LVH, we performed TAC or sham surgery in mice with cardiomyocyte-specific, conditional deletion of *Trpm4* (*Trpm4*^cKO^) using Cre expression driven by the *MLC2a* promoter (42). Results obtained in these *Trpm4*^cKO^ mice were compared with those in WT (*Trpm4^+/+^*) mice. Hemodynamic and anatomical parameters obtained after 2 days and 14 days of sham/TAC in WT and *Trpm4*^cKO^ mice are shown in **Supplementary Table 2**. TAC produced a similar degree of LV pressure overload in both WT (*p* < 0.001) and *Trpm4*^cKO^ (*p* < 0.001) mice when compared with sham-operated groups (**Fig. 4A**), but did not alter HR (**Fig. 4B**), cardiac contractility (**Fig. 4C, D**), BW (**Fig. 4G**) or LW (**Fig. 4E**). **Fig. 4F** illustrates representative images of WT and *Trpm4*^cKO^ mouse hearts after 14 days of sham or TAC. No LVH was detected 2 days after TAC in either *Trpm4*^cKO^ mice or WT mice when compared with sham-operated groups (**Fig. 4G-J**). After 14 days, TAC induced a 32, 42 and 44% increase (all p <0.001) in HW/BW ratio, LVW/BW ratio and LVW/TL ratio, respectively, in WT mice when compared with sham-operated controls (**Fig. 4G-J**). However, this hypertrophic response to 14 days of TAC was attenuated in *Trpm4*^cKO^ mice, as evident by only a 17, 20 and 23% increase (all *p* < 0.001) in HW/BW ratio, LVW/BW ratio and LVW/TL ratio, respectively (**Fig. 4G-J**). These findings demonstrate that when compared with WT mice, *Trpm4*^cKO^ mice developed approximately 50% less LVH (*p* < 0.001) in response to TAC.

**Fig. 4.**
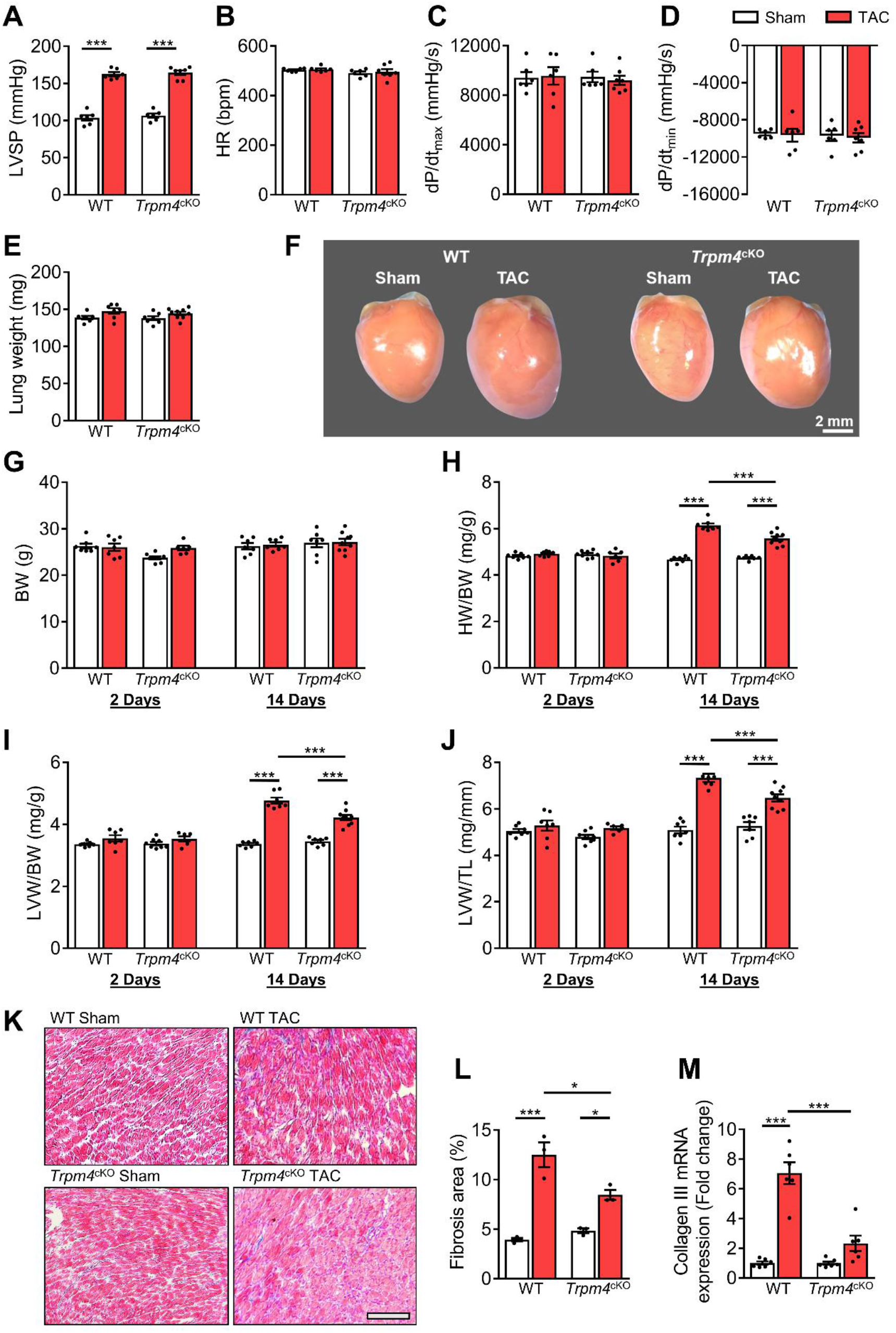
The hypertrophic response of WT and *Trpm4*^cKO^ mice to TAC-induced LV pressure overload. (*A*) Systolic pressure, (*B*) heart rate, (*C*) and (*D*) dP/dt after 14 days of sham or TAC in WT and *Trpm4*^cKO^ mice. (n = 6-7/group); (*E*) Lung weight after 14 days of sham or TAC in WT and *Trpm4*^cKO^ mice. (n = 7-9/group); *(F)* Representative photos indicated heart size differences after 14 days of sham or TAC in WT and *Trpm4*^cKO^ mice; *(G)* Body weight, *(H)* Heart weight, and *(I, J)* LV weight normalized to body weight and tibia length, in WT and *Trpm4*^cKO^ mice after 2 days and 14 days of sham or TAC. (n = 7-9/group); *(K)* Representative micrographs and *(L)* Quantitation of Masson’s trichrome staining of LV tissue from WT mice and *Trpm4*^cKO^ mice after 14 days of sham or TAC by (n = 3/group), scale bar = 200 μm in *(K)*; *(M)* Relative *Col3a1* mRNA expression after 14 days of sham or TAC. (n = 6/group). The mRNA relative expression was calculated as fold change, normalized by GAPDH. Results are presented as means ± SEM. \**p* < 0.05, \*\*\**p* < 0.001.

### Reduced fibrosis in Trpm4^cKO^ hearts after TAC

We evaluated cardiac fibrosis in response to pressure overload in *Trpm4*^cKO^ hearts and WT hearts by Masson’s trichrome staining (**Fig. 4K**). Compared to an average 3.17-fold increase (*p* < 0.001) in cardiac fibrosis in WT TAC hearts, the increase in *Trpm4*^cKO^ TAC hearts was only 1.75-fold (*p* < 0.05) (**Fig. 4L**). In addition, we found a significant increase in collagen III mRNA expression in WT TAC hearts compared to WT sham hearts (*p* < 0.001). However, collagen III mRNA expression was unaltered in *Trpm4*^cKO^ TAC hearts compared to sham hearts (**Fig. 4M**). Thus, *Trpm4* inactivation attenuated the fibrotic response to TAC.

### TRPM4 deficiency reduced the expression of hypertrophy markers in response to TAC-induced pressure overload

Consistent with the development of pathological hypertrophy, both 2 and 14 days of TAC in WT mice significantly enhanced expression of the hypertrophy-associated genes, ANP (*Nppa*), BNP (*Nppb*) and α-SA (*Acta1*) (**Fig. 5A and B**). However, these gene markers remained unchanged with TAC in *Trpm4*^cKO^ mice (**Fig. 5B**), except for ANP (*Nppa*) at 14 days. These data indicate that loss of TRPM4 attenuates the activation of hypertrophic marker genes in response to TAC.

**Fig. 5.**
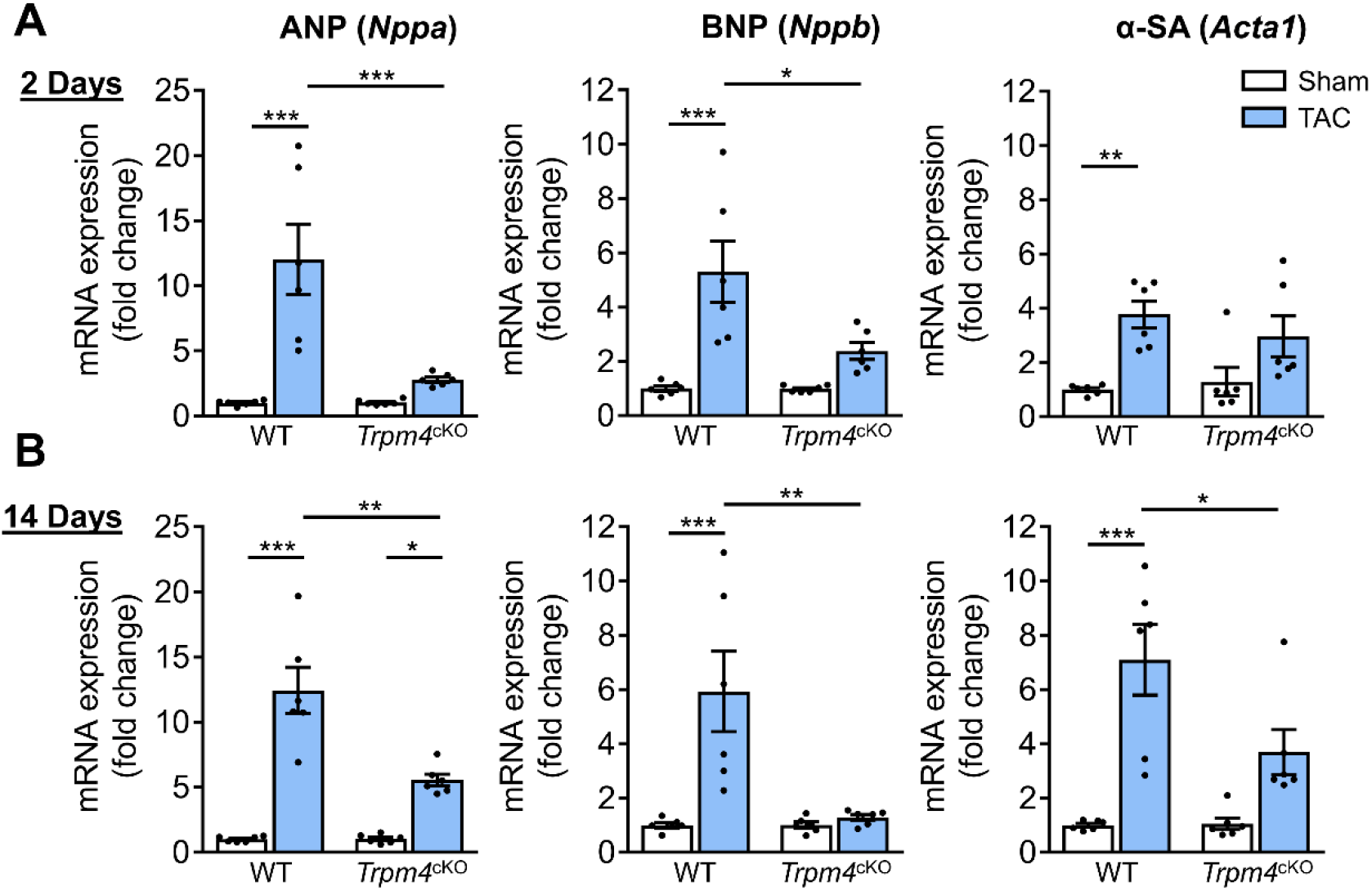
Comparison of gene expression of LVH markers in response to TAC-induced pressure overload in WT and TRPM4^cKO^ mice. *(A)* Relative mRNA expression of ANP (*Nppa*), BNP (*Nppb*) and α-SA (*Acta1*) after 2 days of TAC compared to sham-operated mice. (n = 6/group). *(B)* Relative mRNA expression of ANP (*Nppa*), BNP (*Nppb*) and α-SA (*Acta1*) after 14 days of sham and TAC. (n = 6/group). The mRNA relative expression was calculated as fold change, normalized to GAPDH. Results are presented as means ± SEM, \**p* < 0.05, \*\**p* < 0.01, \*\*\**p* < 0.001.

### CaMKIIδ-HDAC4-MEF2A hypertrophic signalling pathway in WT and Trpm4^cKO^ mouse hearts

We next examined the molecular signalling pathways mediating LVH in both WT and *Trpm4*^cKO^ hearts after 2 days of TAC, a time at which molecular signalling is already activated in response to TAC-induced acute myocardial stretch, but before measurable LVH has developed.

Representative images of key cytoplasmic and nuclear proteins detected by Western blot analysis are shown in **Fig. 6A**. Quantitative data for cytoplasmic and nuclear proteins, normalized by GAPDH and Histone H2B, respectively, are shown in **Fig. 6B**. In WT hearts, 2 days of TAC resulted in a significant increase in CaMKIIδ protein levels in both the cytoplasm (*p* < 0.01) and nucleus. Associated with this increase, there was a rise in total cytoplasmic HDAC4 (*p* < 0.01) and phosphorylated HDAC4 (p-HDAC4) levels (*p* < 0.001), but no change in nuclear HDAC4. This 2.11-fold increase in the cytoplasmic/nuclear ratio of HDAC4 (*p* < 0.01) in WT hearts indicates that TAC-induced pressure overload leads to the nuclear export of HDAC4 in WT TAC hearts. This increase was accompanied by a 1.76-fold increase of MEF2A levels in the nucleus (*p* < 0.05), which together with the de-repression of MEF2A activity would account for the induction of LVH.

**Fig. 6.**
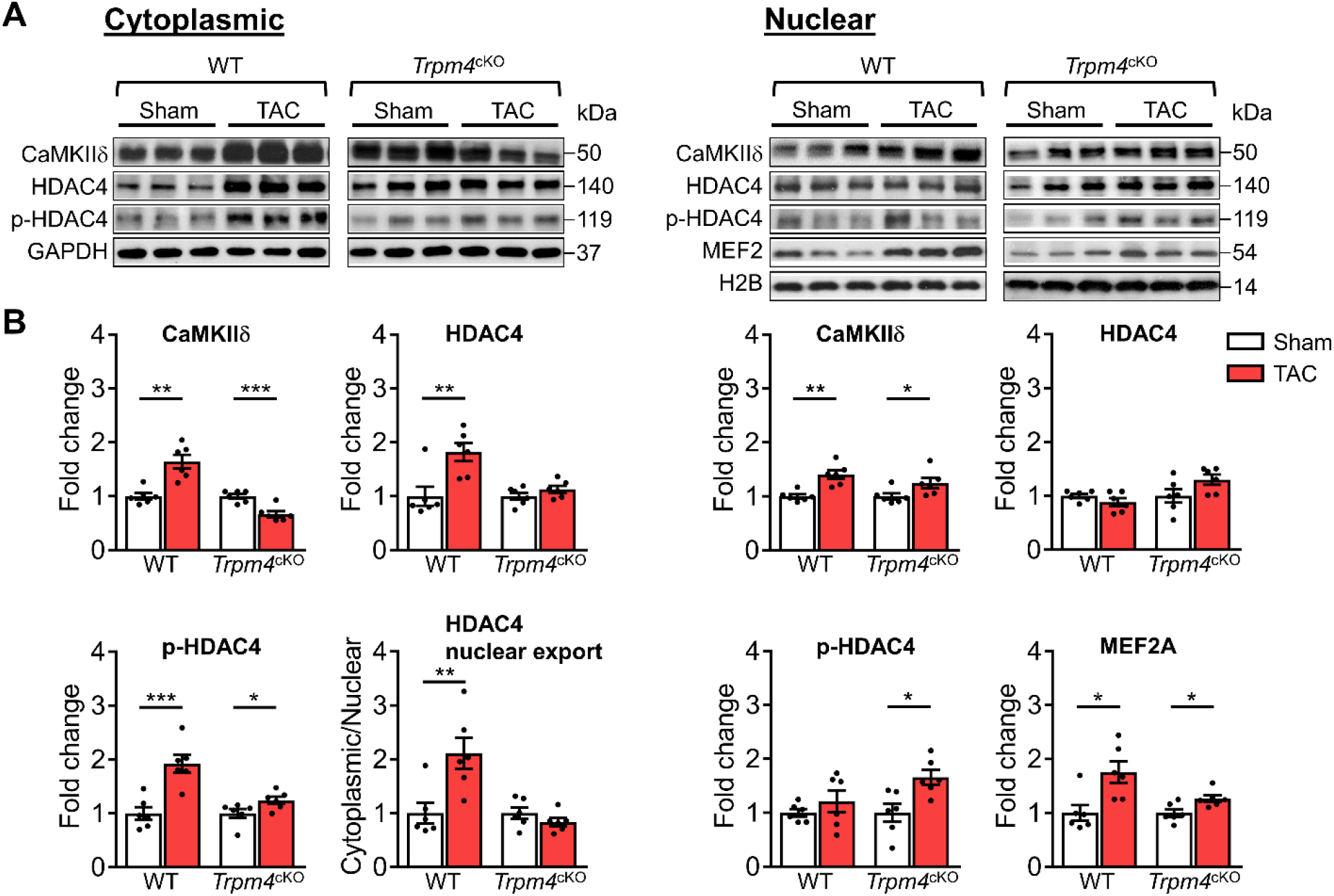
CaMKIIδ, HDAC4 and MEF2A signalling pathway in response to TAC after 2 days in WT and *Trpm4*^cKO^ mouse hearts. (*A*) Representative western blots showing the change of key proteins in this signalling pathway in the cytoplasm (*left panel*) and nucleus (*right panel*). (*B*) Cytoplasmic (*left panel*) and nuclear (*right panel*) quantitative data were normalized by GAPDH and Histone H2B respectively. HDAC4 nuclear export was determined using the HDAC4 cytoplasmic/nuclear ratio. Results are presented as means ± SEM, n = 6/group, \**p* < 0.05, \*\**p* < 0.01, \*\*\**p* < 0.001.

**Fig. 7.**
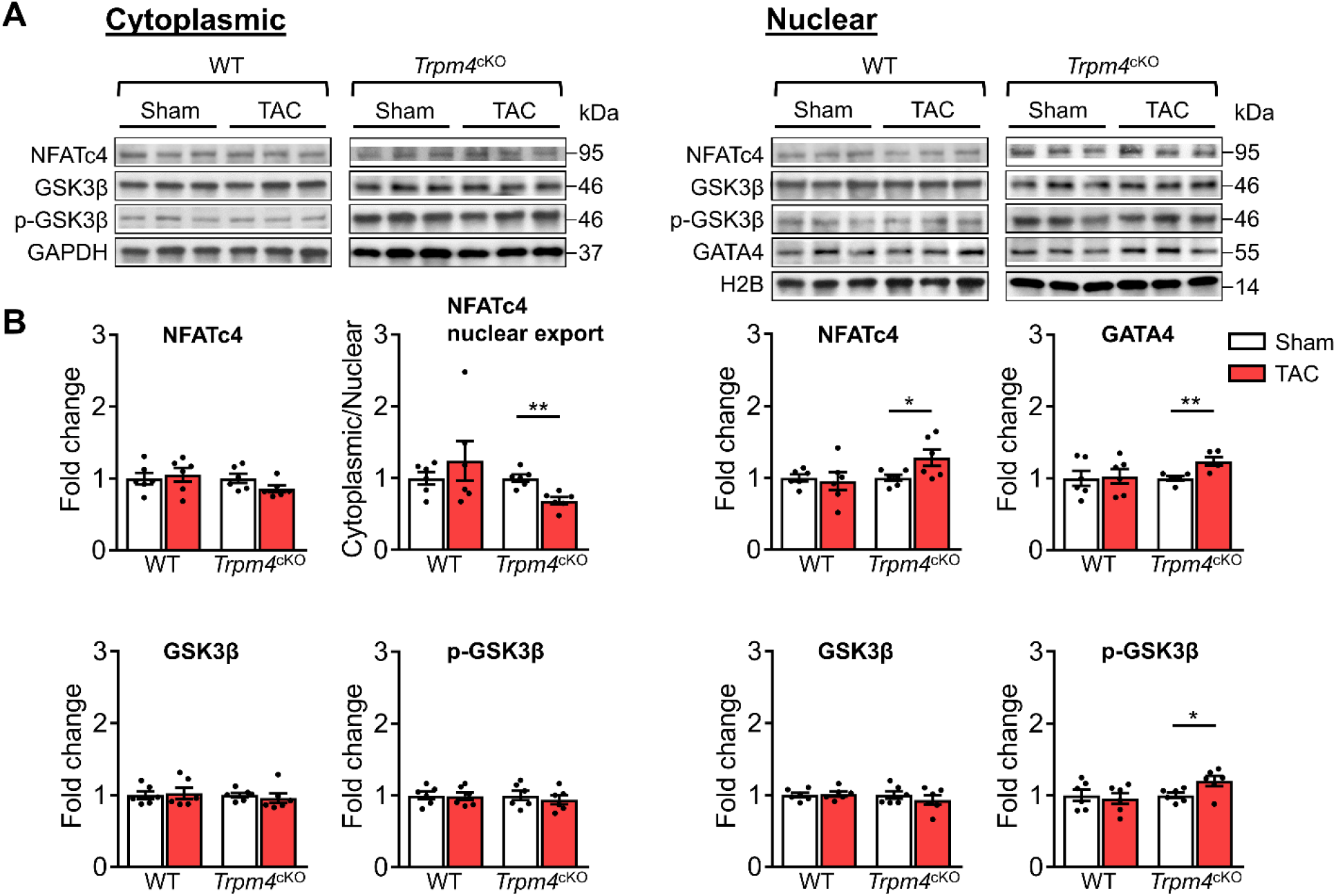
Calcineurin-NFAT-GATA4 signalling pathway in response to TAC after 2 days in WT and *Trpm4*^cKO^ mouse hearts. (*A*) Representative western blots showing the change of key proteins in the calcineurin-NFAT-GATA4 signalling pathway in cytoplasm (*left panel*) and nucleus (*right panel*). (*B*) Cytoplasmic (*left panel*) and nuclear (*right panel*) quantitative data were normalized by GAPDH and Histone H2B respectively. NFATc4 nuclear export was indicated with its cytoplasmic/nuclear ratio. Results are presented as means ± SEM, n = 6/group, \**p* < 0.05, \*\**p* < 0.01.

**Fig. 8.**
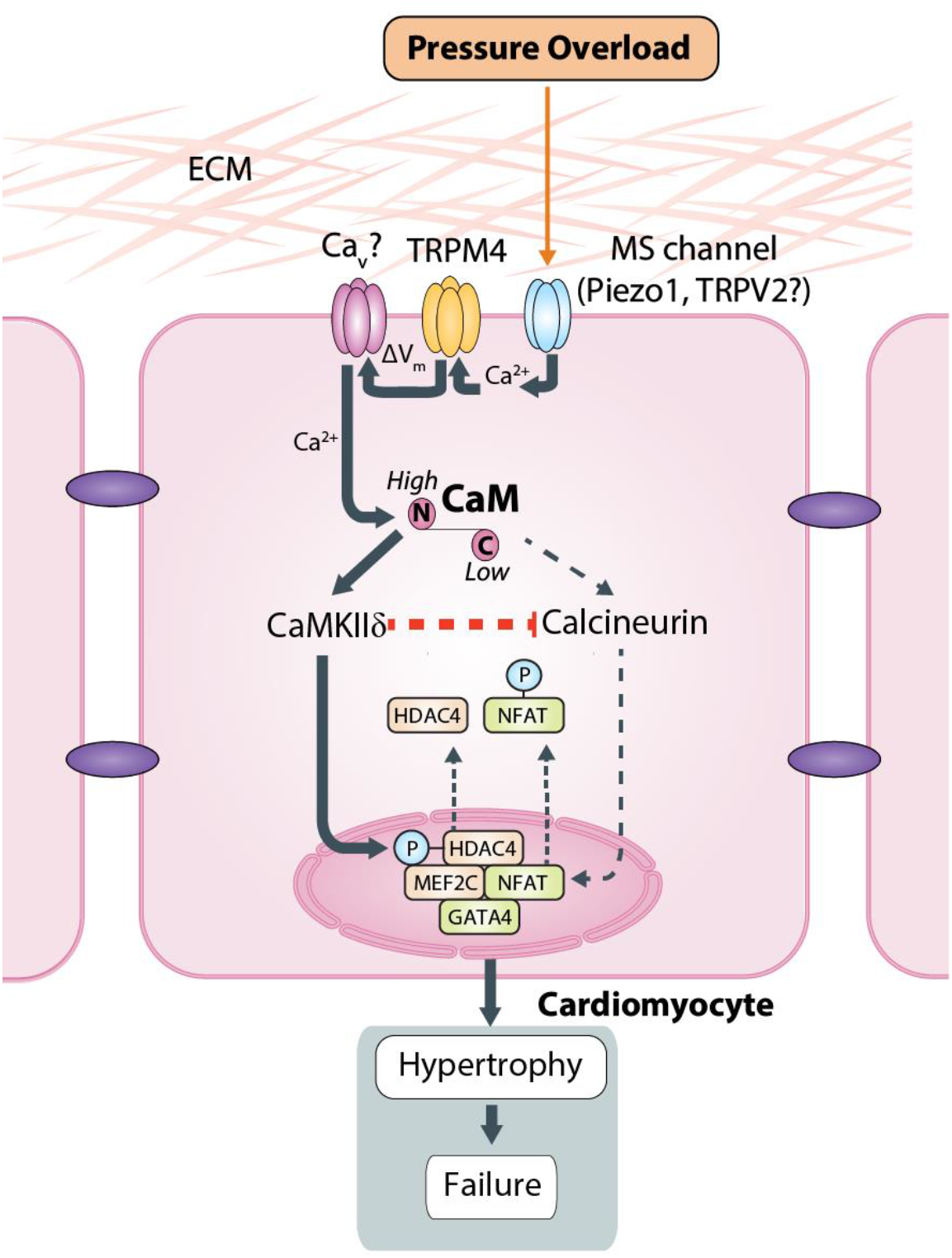
Schematic of the putative TAC-induced pathway that culminates in LVH. CaM - calmodulin; Ca_V_ - voltage gated Ca^2+^ channel.

In contrast to the effects of TAC in WT hearts, in *Trpm4*^cKO^ hearts, TAC produced a lesser increase (0.66-fold of that observed in sham hearts; *p* < 0.001) in cytoplasmic CaMKIIδ levels, whereas the increase in nuclear CaMKIIδ was similar (*p* < 0.05) to that observed with TAC in WT hearts. p-HDAC4 also increased in TAC *Trpm4*^cKO^ hearts in both the cytoplasm (*p* < 0.05) and the nucleus (*p* < 0.05) but there was no change in total HDAC4. Thus, the cytoplasmic/nuclear ratio of HDAC4 remained the same in *Trpm4*^cKO^ hearts as in sham hearts, indicating inhibition of nuclear HDAC4 export in TAC-treated *Trpm4*^cKO^ hearts. In addition, consistent with MEF2A activation driving hypertrophy development, reduced LVH in the TAC *Trpm4*^cKO^ hearts was associated with a lesser (1.26-fold) increase in MEF2A concentration in the nucleus (*p* < 0.05). Taken together, these data implicate the CaMKIIδ-HDAC4-MEF2A hypertrophic signalling pathway in mediating TAC-induced LVH, the extent of which is regulated by TRPM4 channels.

### Calcineurin-NFAT-GATA4 hypertrophic signalling pathway in WT and Trpm4^cKO^ mouse hearts

Next, we examined the expression of proteins involved in the calcineurin-NFAT-GATA4 hypertrophic signalling pathway. In WT hearts, there was no significant difference in cytoplasmic or nuclear NFATc4 protein expression in sham and TAC hearts after 2 days. Consistent with these findings, total GSK3β, serine-9 phosphorylated GSK3β and GATA4 levels were also unchanged in response to TAC, confirming our previous finding (23) that the calcineurin-NFAT-GATA4 pathway is not activated by TAC.

In contrast to WT hearts, an increase in nuclear NFATc4 (*p* < 0.05) was observed in *Trpm4*^cKO^ hearts after TAC, which led to a 0.69-fold decrease in the cytoplasmic/nuclear ratio compared to sham-operated hearts (*p* < 0.01). This indicated lower nuclear export of NFATc4 in the *Trpm4*^cKO^ TAC hearts when compared to sham hearts. Accordingly, we found a 1.20-fold increase in nuclear p-GSK3β (ser-9) (*p* < 0.05) in *Trpm4*^cKO^ TAC hearts. As phosphorylation at the serine-9 residue indicates inactivation of GSK3β, these findings suggest that the GSK3β-mediated export of NFATc4 from the nucleus was partially inhibited, which is consistent with the increased level of NFATc4 in the nucleus. Furthermore, accompanied by the increase in nuclear NFATc4, a 1.18-fold increase in GATA4 expression (*p*< 0.05) in the nucleus was observed in *Trpm4*^cKO^ TAC hearts. All these observations are consistent with a reduction in the tonic inhibition of calcineurin by CaMKIIδ (55) in *Trpm4*^cKO^ hearts after TAC.

## Discussion

In the present study, we employed mice subjected to TAC as an *in vivo* disease model to investigate the role of the TRPM4 ion channels in pressure overload-induced pathological LVH. The experimental animals were examined 2 days after surgery when the molecular signalling pathway that drives LVH is switched on in response to the acute stretch generated by TAC but, importantly, before LVH has developed. In addition, we examined animals from the different groups at 14 days after surgery when the TAC-induced LVH phenotype is evident.

First, we found that TRPM4 channel expression in the WT mouse heart was modified by TAC-induced pressure overload hypertrophy. At 2 days and 14 days after TAC, both *Trpm4* mRNA and protein expression were downregulated in LV tissue and isolated cardiomyocytes, suggesting that TRPM4 plays a pro-hypertrophic role in TAC-induced LVH. Second, we confirmed this pro-hypertrophic role of TRPM4 by performing sham and TAC surgery in *Trpm4*^cKO^ mice. This demonstrated that a reduction in TRPM4 expression in cardiomyocytes dampens the hypertrophic response to TAC, as evident by an approximately 50% reduction in the degree of LVH and LV fibrosis in *Trpm4*^cKO^ animals at 14 days after TAC, as compared to WT animals. Finally, to investigate the hypertrophic signalling pathways activated in response to pressure overload, we examined both the CaMKIIδ-HDAC4-MEF2A and calcineurin-NFAT-GATA4 signalling pathways 2 days after TAC in WT and *Trpm4*^cKO^ mice (**Fig. 8**). This revealed reduced activation of the CaMKIIδ-HDAC4-MEF2A pathway 2 days in *Trpm4*^cKO^ animals.

There is evidence that the TRPM4 channel is a critical modulator of ventricular remodelling in cardiac hypertrophy and heart failure (38–40, 42). Previous studies suggest that TRPM4 activation supresses angiotensin II-induced cardiac hypertrophy, dependent on the activation of the calcineurin-NFAT pathway. This is due to the Ca^2+^-dependent modulation of TRPM4 activity, which leads to membrane depolarization in cardiomyocytes and thus reduces the driving force for Ca^2+^ influx via store-operated calcium entry (SOCE) through TRPC1 and TRPC3 ion channels (42, 43). However, to our knowledge, a role for TRPM4 in the LVH induced by mechanical pressure overload has not been demonstrated previously. We propose here that a mechanical stimulus, such as that exerted by TAC, is converted to downstream Ca^2+^ signaling via the activity of mechanosensitive ion channels in the plasma membrane. Although the mechanosensitivity of TRP-type ion channels is still the subject of debate (44, 45), mammalian TRP ion channels, including TRPM4, have recently been shown to be insensitive to membrane stretch (46, 47). Therefore, TRPM4 does not appear to be the primary mechanosensor responding to pressure overload. It is more likely to be a secondary ionotropic receptor downstream of a calcium-permeable mechanosensitive ion channel, such as Piezo1 (48) or TRPV2/4 (49), the latter functioning as the primary mechanoreceptor responding directly to pressure overload and thus initiating the hypertrophic response in TAC.

Influx of monovalent cations (e.g. Na^+^) through TRPM4 would depolarize the cardiomyocyte cell membrane, which could activate voltage-gated Ca^2+^ channels allowing further entry of extracellular calcium. Thus, as a Ca^2+^-dependent non-selective monovalent cation channel (33, 46, 50, 51), TRPM4 could contribute to TAC-induced LVH by modulating downstream voltage-gated Ca^2+^ ion channels (**Fig. 8**). Such potential downstream ion channels include L-type Ca^2+^ channels, which were reported to mediate hypertrophic cardiomyopathy (52) as well as T-type Ca^2+^ channels, whose splice variants were found to be regulated in the LV of rat hearts made hypertrophic by aortic constriction (53).

We confirmed the involvement of TRPM4 in TAC-induced LVH using *Trpm4*^cKO^ mice. Despite identical TAC-induced increases in hemodynamic load in both WT and *Trpm4*^cKO^ mice, the latter displayed a significantly reduced LVH response. This is in contrast to the increased hypertrophy reported in angiotensin II treated *Trpm4*^cKO^ mice that is mediated by the calcineurin-NFAT pathway (42). These differential effects of TRPM4 on angiotensin II-mediated (42) and TAC-induced LVH support our previous finding that these two hypertrophic stimuli are mediated by distinct signalling mechanisms. Thus, in agreement with the present findings, we showed that the CaMKIIδ-HDAC4-MEF2A, but not the calcineurin-NFAT-GATA4 signalling pathway, is activated in response to TAC-induced pressure overload (23). This is most likely because CaMKII and calcineurin respond to different characteristics of intracellular Ca^2+^ signaling (54, 55). Whereas calcineurin activation requires a sustained increase in the resting intracellular Ca^2+^ concentration, CaMKII activation is more sensitive to high-frequency/high amplitude calcium oscillations (55, 56), which are known to occur with TAC-induced aortic constriction (57). Consequently, a possible mechano-transduction scenario in TAC-induced pressure overload is activation of mechanosensitive Ca^2+^-permeable ion channels, such as Piezo1 (48) or TRPV2/4 (49), as the load-dependent source of extracellular calcium; the resulting increase in intracellular Ca^2+^ activating TRPM4 channels.

In terms of the signalling pathway mediating pressure overload-induced LVH, we found that in *Trpm4*^cKO^ TAC hearts, the reduced LVH response was associated with significantly less activation of the CaMKIIδ-HDAC4-MEF2A pathway; reduced CaMKIIδ activation resulting in reduced nuclear export of HDAC4. Since nuclear HDAC4 inhibits MEF2A activity, a reduction in HDAC4 nuclear export would result in diminished MEF2A disinhibition and, given that MEF2A is a critical nuclear transcriptional regulator causing pathological cardiac remodelling, reduced hypertrophy development (25) as, indeed, observed here in *Trpm4*^cKO^ TAC hearts.

Decreased expression of TRPM4 channels in *Trpm4*^cKO^ animals likely modifies Ca^2+^-signaling, which directly regulates CaMKIIδ activation and its downstream pathway in response to TAC. Comparable with a study reporting that blockade of MEF2 acetylation can permit recovery from pathological cardiac hypertrophy without impairing physiologic adaptation (58), the lower concentration and reduced activity of MEF2A that we found in *Trpm4*^cKO^ TAC hearts suggest that inhibition of TRPM4 channels is potentially a viable therapeutic option for reducing pathological hypertrophic in response to pressure overload.

Interestingly, although the calcineurin-NFAT-GATA4 hypertrophic signalling pathway is not activated by TAC in WT hearts, it was partially activated in *Trpm4*^cKO^ TAC hearts, which manifested itself in the inhibition of GSK3β-mediated NFATc4 nuclear export and by an increase in GATA4. This may be explained by the reduction of the cytoplasmic CaMKIIδ in *Trpm4*^cKO^ TAC hearts, as CaMKII has been reported to negatively regulate calcineurin activity (59). It is notable, nevertheless, that the net effect of the loss of TRPM4 was a significant reduction in TAC-induced LVH, indicating that the direct effect of less activation of the CaMKIIδ-HDAC4-MEF2A pathway is reduced hypertrophy development, which outweighs the indirect effects resulting from blunting CaMKIIδ’s inhibition of calcineurin.

In summary, our study provides compelling evidence that TRPM4 plays an important role in pressure overload-induced pathological LVH, with diminished TRPM4 expression reducing TAC-induced hypertrophy. Furthermore, we demonstrated that TRPM4 is a likely component of a cardiac mechano-transduction process that activates the CaMKIIδ-HDAC4-MEF2A pathway in response to TAC. It is likely that TRPM4 is activated by upstream primary mechanoreceptors, such as Piezo1 or TRPV2/4 channels, which provide the first step in this mechano-transduction pathway. These findings expand our understanding of the molecular mechanism underlying mechanical pressure-overload-induced LVH. Moreover, our work provides new insights into possible treatment strategies for limiting pressure overload-induced pathological hypertrophy.

## Materials and Methods

### Mice

In the first part of the study, we performed experiments on 11 to 13-week-old male C57BL/6J WT mice at the Victor Chang Cardiac Research Institute, Australia. In the second part of this study, we performed surgery on C57BL/6N WT and age-and sex-matched cardiac-specific *Trpm4*^cKO^ mice in KU Leuven, Belgium. *Trpm4*^flox^ mice were crossbred with MLC2a-Cre mice to generate the *Trpm4*^cKO^ mice (42). All animals were entered into the study in a randomized order, and the investigators were blinded to genotype. All experimental procedures were approved by the Animal Ethics Committee of Garvan/St Vincent’s (Australia) or KU Leuven (Belgium), respectively, in accordance with the guidelines of both the Australian code for the care and use of animals for scientific purposes (8th edition, National Health and Medical Research Council, AU, 2013) and the Guide for the Care and Use of Laboratory Animals (8th edition, National Research Council, USA, 2011).

### Induction of LVH

WT and *Trpm4*^cKO^ mice were subjected to TAC to induce pressure overload. Mice were anesthetized with 5% isoflurane and ventilated at 120 breaths/min (Harvard Apparatus Rodent Ventilator). The transverse aortic arch was accessed via an incision in the second intercostal space, and constricted with a ligature tied around a 25-gauge needle, which was then removed. The TAC procedure was modified from a published paper (60). Sham mice underwent the same procedure but the ligature was not tied. Simultaneous direct pressure recordings (1.4 F pressure catheter, AD Instruments, P/L) from both the right carotid artery and the aorta distal to the ligature (n=20 mice) indicated a TAC pressure gradient of 60 ± 8 mmHg with this technique. Animals were sacrificed after 2 days or 14 days.

### Invasive hemodynamic measurements

After 14 days of sham or TAC, mice were anesthetized by inhalation of isoflurane (1.5%) and a 1.4F micro-tip pressure catheter (Millar Instruments Inc, Houston, Texas, USA) was inserted into the left ventricle via the right carotid artery. The heart rate, systolic aortic pressure, LV systolic pressure, +dP/dt, and –dP/dt were recorded (LabChart 6 Reader, AD Instruments, P/L). Animals were sacrificed, and the heart weight (HW) and left ventricle weight (LVW) normalized to body weight (BW) and to tibia length (TL) were measured as indicators of LVH.

### Mouse LV cardiomyocytes isolation and purification

WT mice were heparinized and euthanized according to the Animal Research Act 1985 No 123 (New South Wales, Australia). Hearts were dissected and perfused through the aorta and the coronary arteries by 10 ml pH 7.2 perfusion buffer containing 135 mM NaCl, 4 mM KCl, 1 mM MgCl_2_, 0.33 mM NaH_2_PO_2_, 10 mM HEPES, 10 mM Glucose, 10 mM 2,3-Butanedione 2-monoxime (BDM), and 5 mM Taurine, with a Langendorff apparatus at 37 degrees for 5 minutes. Next, 30 ml digestion buffer composed of the above solution and Collagenase B, D (dose by BW: 0.4 mg/g, Roche) and Protease Enzyme Type XIV (dose by BW: 0.07 mg/g, Sigma-Aldrich) was used to perfuse the hearts for 15 minutes. After the perfusion, the heart was removed from the setup and placed into a pH 7.4 transfer buffer containing 135 mM NaCl, 4 mM KCl, 1 mM MgCl_2_, 0.33 mM NaH_2_PO_2_, 10 mM HEPES, 5.5 mM Glucose, 10 mM BDM, and 5 mg/ml BSA. Both atria and the right ventricle were discarded, and the LV muscle was torn into small pieces and gently dispensed into the transfer buffer repeatly with a pipette to isolate cardiomyocytes. The suspension was then filtered through a 200 micro filcon cup filter (BD), and centrifuged at 20 g for 2 minutes. After that, the cardiomyocytes were purified by a method that provides 95% purity using Purified Rat Anti-Mouse CD31 antibody (BD Pharmingen) and Dynabeads Sheep Anti-Rat IgG (Invitrogen), which will be detailed in a separate paper (61). We confirmed that rod-shaped cardiomyocytes accounted for more than 85% of the total purified cardiomyocytes. The isolated cardiomyocytes were frozen immediately in liquid nitrogen and stored at −80 degrees for following experiments.

### Quantitative Real-Time Polymerase Chain Reaction (RT-PCR)

Gene expression was determined by quantitative RT-PCR. Total RNA was extracted and purified from LV tissue and isolated cardiomyocytes with the RNeasy Fibrous Tissue Mini Kit (QIAGEN), following the manufacturer’s protocol. RNA (500 ng) was reverse transcribed into cDNA using the SuperScript III First-Strand Synthesis SuperMix kit (Invitrogen). cDNA was subjected to PCR amplification to detect ANP (*Nppa*), BNP (*Nppa*), α-SA (*Acta1*), collagen 3 (*Col3a1*), and *Trpm4* gene expression, performed with the CFX384 Touch Real-Time PCR Detection System (Bio-Rad), PCR master mix LightCycler 480 SYBR Green I Master (Invitrogen). Samples were run in technical triplicate and the mRNA expression levels were normalized to those of GAPDH to calculate relative gene expression using delta-delta Ct method. The mouse RT-PCR primers (Sigma-Aldrich) used are shown in **Supplementary Tab. 3**.

### Western blotting

For total protein extraction, LV tissue and isolated cardiomyocytes were lysed in a pH 7.4 lysis buffer containing 150 mM NaCl, 50 mM Tris-HCL, 1% Triton X-100, 1 mM sodium orthovanadate, 1 mM beta-glycerophosphate, 5 mM Dithiothreitol, and MiniComplete protease inhibitors (Roche); for cytoplasmic and nuclear protein extraction, LV tissue was lysed using NE-PER nuclear and cytoplasmic extraction reagents (Pierce Biotechnology) and Protesase Inhibitor Cocktail Kit and Halt Phosphatase Inhibitor Cocktail (Pierce Biotechnology), both with a homogenizer (PRO Scientific). Protein (40 μg for each sample) was loaded on 4%-20% Mini-PROTEAN TGX Gels (Bio-Rad) and separated by electrophoresis. Samples were transferred to PVDF membranes (Bio-Rad), blocked with 5% bovine serum albumin (BSA) then labelled overnight with primary antibodies: anti-TRPM4 (1:200, Alomone Labs), anti-CaMKIIδ (1:1000; Santa Cruz Biotechnology), anti-total HDAC4 (1:1500; Cell Signalling), anti-p-HDAC4 (1:1500; Cell Signalling), anti-MEF2A (1:3000; Cell Signalling), anti-NFATc4 (1:1500 final dilution; Abcam), anti-total GSK3β (1:500; Cell Signalling), anti-p-GSK3β (Ser9, 1:1500; Cell Signalling), anti-GATA4 (1:1000; Santa Cruz Biotechnology). Anti-Glyceraldehyde-3-phosphate dehydrogenase (GAPDH, 1:5000; Abcam) and anti-Histone H2B (1:5000; Abcam) were used to standardize for loading. Horseradish peroxidase-conjugated goat anti-mouse (1:5000) or anti-rabbit (1:10000) secondary antibodies (Abcam) were used at room temperature for one hour. Immunologic detection was accomplished using Amersham ECL Western blotting detection reagents (GE Healthcare). Protein levels were quantified by densitometry using ImageJ (NIH) software. Protein levels were normalized to relative changes in Histone H2B for the nuclear fraction and GAPDH for the cytoplasmic fraction and expressed as fold changes relative to those of control animals.

### Histology

Dissected mouse hearts from both WT and *Trpm4*^cKO^ groups were perfused with phosphate-buffered saline (PBS). Then the hearts were embedded into optimal cutting temperature (OCT) compound (Sakura Finetek), gradually frozen in liquid nitrogen via isopentane to avoid tissue damage. Serial sections with a thickness of 6 microns were sliced with a cryostat (Leica). The slides were then stained with a masson’s trichrome staining kit (Sigma-Aldrich) following the manufacturer’s instructions, and imaged with a brightfield microscope (Leica). The obtained images were quantified by ImageJ (NIH).

### Statistics

All experiments and analyses were blinded. Averaged data are presented as means ± standard error of the mean (SEM). The statistical analysis was performed using GraphPad Prism software, version 7.04 (GraphPad). For comparisons between two sets of data, unpaired t-test was used to determine the statistical significance. For comparisons among multiple sets of data with one factor or two factors, one-way or two-way ANOVA was used accordingly, followed by Tukey’s post-hoc test. *p* < 0.05 was considered statistically significant.

## Acknowledgements

The authors gratefully acknowledge funding from the National Health and Medical Research Council (NHMRC) of Australia to BM and MF through a project grant (APP1108013), as well as the NHMRC of Australia for a Principal Research Fellowship (APP1135974) and NSW Cardiovascular Disease Senior Scientist Grant to BM. RV, SP and AP are supported by the BOF KU Leuven (TRPLe) and the FWO-Vlaanderen (G0E0317N). CDC is supported by an NSW Health EMCR Fellowship. This work is part of a PhD thesis of YG.

---

**Supplementary Tab. 1.** Hemodynamic and anatomical parameters 2 days or 14 days after induction of pressure overload in WT mice.

|  | 2 days |  | 14 days |  |
| --- | --- | --- | --- | --- |
|  | Sham | TAC | Sham | TAC |
| **Hemodynamic parameter** |  |  |  |  |
| n |  |  | 7 | 7 |
| HR (bpm) |  |  | 503 ± 7 | 507 ± 6 |
| Aortic systolic pressure (mmHg) |  |  | 103 ± 3 | 166 ± 9 *** |
| Aortic diastolic pressure (mmHg) |  |  | 76 ± 1.5 | 74 ± 1.8 |
| LV systolic Pressure (mmHg) |  |  | 105 ± 2.8 | 164 ± 8 *** |
| dP/dt<sub>max</sub> (mmHg/s) |  |  | 9449 ± 220 | 9838 ± 250 |
| dP/dt<sub>min</sub> (mmHg/s) |  |  | -9666 ± 218 | -9998 ± 240 |
| **Anatomical parameter** |  |  |  |  |
| n | 8 | 8 | 11 | 11 |
| BW (g) | 28.5 ± 0.8 | 27.7 ± 1 | 28.6 ± 0.9 | 27.5 ± 1.1 |
| HW (mg) | 136 ± 2 | 133 ± 1 | 131 ± 2 | 176 ± 8 *** |
| LVW (mg) | 98 ± 1.8 | 97.6 ± 1.2 | 96 ± 1 | 136 ± 4 *** |
| LW (mg) | 141 ± 1 | 143 ± 1.5 | 146 ± 1.8 | 147 ± 2 |
| TL (mm) | 17.4 ± 0.8 | 17.3 ± 1 | 17.4 ± 1.0 | 17.3 ± 0.9 |
| HW/BW (mg/g) | 4.8 ± 0.1 | 4.75 ± 0.1 | 4.68 ± 0.1 | 6.6 ± 0.2 *** |
| LVW/BW (mg/g) | 3.4 ± 0.07 | 3.5 ± 1 | 3.37 ± 0.08 | 5.09 ± 0.1 *** |
| LVW/TL (mg/mm) | 5.6 ± 0.2 | 5.7 ± 0.2 | 5.54 ± 0.1 | 7.9 ± 0.15 |
| LW/BW (mg/g) | 5.4 ± 0.2 | 5.3 ± 0.1 | 5.5 ± 0.09 | 5.4 ± 0.1 |

HR, heart rate. LV, left ventricle. dP/dt<sub>max</sub>, dP/dt<sub>min</sub>, BW, body weight. LVW, left ventricular weight. LW, lung weight. TL, tibia length. HW/BW, heart weight to body weight ratio. LVW/BW, left ventricular weight to body weight ratio. LVW/TL, left ventricular weight to tibia length ratio. LW/BW, lung weight to body weight ratio. Data expressed as means ± SEM. Comparison between sham and TAC groups: \*\*\**p* < 0.001.

**Supplementary Tab. 2.** Hemodynamic and anatomical parameters after 2 days and 14 days of sham/TAC in WT and *Trpm4*^cKO^ mice.

|  | 2 days |  |  |  | 14 days |  |  |  |
| --- | --- | --- | --- | --- | --- | --- | --- | --- |
|  | WT |  | *Trpm4*<sup>cko</sup> |  | WT |  | *Trpm4*<sup>cko</sup> |  |
|  | Sham | TAC | Sham | TAC | Sham | TAC | Sham | TAC |
| **Hemodynamic parameter** |  |  |  |  |  |  |  |  |
| n |  |  |  |  | 6 | 6 | 6 | 7 |
| HR (bpm) |  |  |  |  | 503.00<br>± 2.78 | 505.83<br>± 4.27 | 491.17<br>± 6.00 | 496.00<br>± 10.67 |
| Aortic systolic pressure (mmHg) |  |  |  |  | 103.83<br>± 2.27 | 161.00<br>± 2.57 *** | 100.83<br>± 3.61 | 163.43<br>± 2.23 *** |
| Aortic diastolic pressure (mmHg) |  |  |  |  | 73.33<br>± 6.31 | 78.67<br>± 2.72 | 72.50<br>± 1.18 | 77.14<br>± 2.72 |
| LV systolic pressure (mmHg) |  |  |  |  | 103.67<br>± 3.68 | 162.50<br>± 2.86 *** | 106.67<br>± 2.86 | 164.43<br>± 3.04 *** |
| dP/dt<sub>max</sub> (mmHg/s) |  |  |  |  | 9403.00<br>± 466.66 | 9559.67<br>± 703.60 | 9470.67<br>± 424.55 | 9199.14<br>± 372.03 |
| dP/dt<sub>min</sub> (mmHg/s) |  |  |  |  | -9492.83<br>± 186.90 | -9642.83<br>± 681.67 | -9706.33<br>± 551.89 | -9924.57<br>± 506.24 |
| **Anatomical parameter** |  |  |  |  |  |  |  |  |
| n | 7 | 7 | 8 | 6 | 7 | 7 | 7 | 9 |
| BW (g) | 26.29<br>± 0.54 | 26.03<br>± 0.80 | 23.75<br>± 0.33 | 25.85<br>± 0.51 | 26.26<br>± 0.67 | 26.60<br>± 0.46 | 26.96<br>± 0.98 | 27.14<br>± 0.69 |
| HW (mg) | 126.57<br>± 1.82 | 128.00<br>± 4.68 | 116.25<br>± 1.77 | 124.50<br>± 1.23 | 122.71<br>± 3.46 | 163.29<br>± 3.79 *** | 127.71<br>± 4.15 | 151.22<br>± 4.52 ** |
| LVW (mg) | 88.29<br>± 1.89 | 92.57<br>± 4.78 | 80.25<br>± 1.26 | 91.00<br>± 1.13 | 88.43<br>± 2.87 | 126.71<br>± 2.96 *** | 93.00<br>± 3.57 | 114.33 #<br>± 2.92 *** |
| LW (mg) | 137.44<br>± 0.98 | 136.63<br>± 2.19 | 130.01<br>± 2.06 | 135.07<br>± 1.69 | 138.66<br>± 2.26 | 147.66<br>± 3.65 | 137.94<br>± 2.75 | 144.38<br>± 2.30 |
| TL (mm) | 17.47<br>± 0.10 | 17.49<br>± 0.20 | 16.63<br>± 0.06 | 17.58<br>± 0.15 | 17.37<br>± 0.09 | 17.29<br>± 0.09 | 17.64<br>± 0.16 | 17.66<br>± 0.09 |
| HW/BW (mg/g) | 4.82<br>± 0.04 | 4.91<br>± 0.04 | 4.90<br>± 0.05 | 4.83<br>± 0.10 | 4.67<br>± 0.04 | 6.14<br>± 0.09 *** | 4.74<br>± 0.03 | 5.57 ###<br>± 0.09 *** |
| LVW/BW (mg/g) | 3.40<br>± 0.03 | 3.55<br>± 0.10 | 3.38<br>± 0.05 | 3.53<br>± 0.08 | 3.36<br>± 0.03 | 4.77<br>± 0.10 *** | 3.45<br>± 0.05 | 4.22 ###<br>± 0.09 *** |
| LVW/TL (mg/mm) | 5.05<br>± 0.09 | 5.28<br>± 0.22 | 4.82<br>± 0.10 | 5.18<br>± 0.07 | 5.09<br>± 0.15 | 7.33<br>± 0.18 *** | 5.27<br>± 0.17 | 6.47 ###<br>± 0.15 *** |
| LW/BW (mg/g) | 5.24<br>± 0.08 | 5.27<br>± 0.09 | 5.48<br>± 0.07 | 5.24<br>± 0.13 | 5.29<br>± 0.09 | 5.55<br>± 0.12 | 5.15<br>± 0.17 | 5.33<br>± 0.09 |

Hemodynamic measurements include heart rate (HR), aortic systolic and diastolic pressure, LV systolic pressure, dP/dt<sub>max</sub> and dP/dt<sub>min</sub>; anatomical measurements include body weight
(BW), heart weight (HW), LV weight (LVW), lung weight (LW), tibial length (TL), heart weight normalized by body weight (HW/BW), LV weight normalized by body weight and tibial length (LVW/BW; LVW/TL), lung weight normalized by body weight (LW/BW). Data are presented as means ± SEM. Comparison between sham and TAC in WT or *Trpm4*<sup>ckO</sup> groups: \*\**p* < 0.01, \*\*\**p* < 0.001; Comparison between WT and *Trpm4*<sup>ckO</sup> TAC groups: #*p* < 0.05, ###*p* < 0.001.

**Supplementary Tab. 3.**
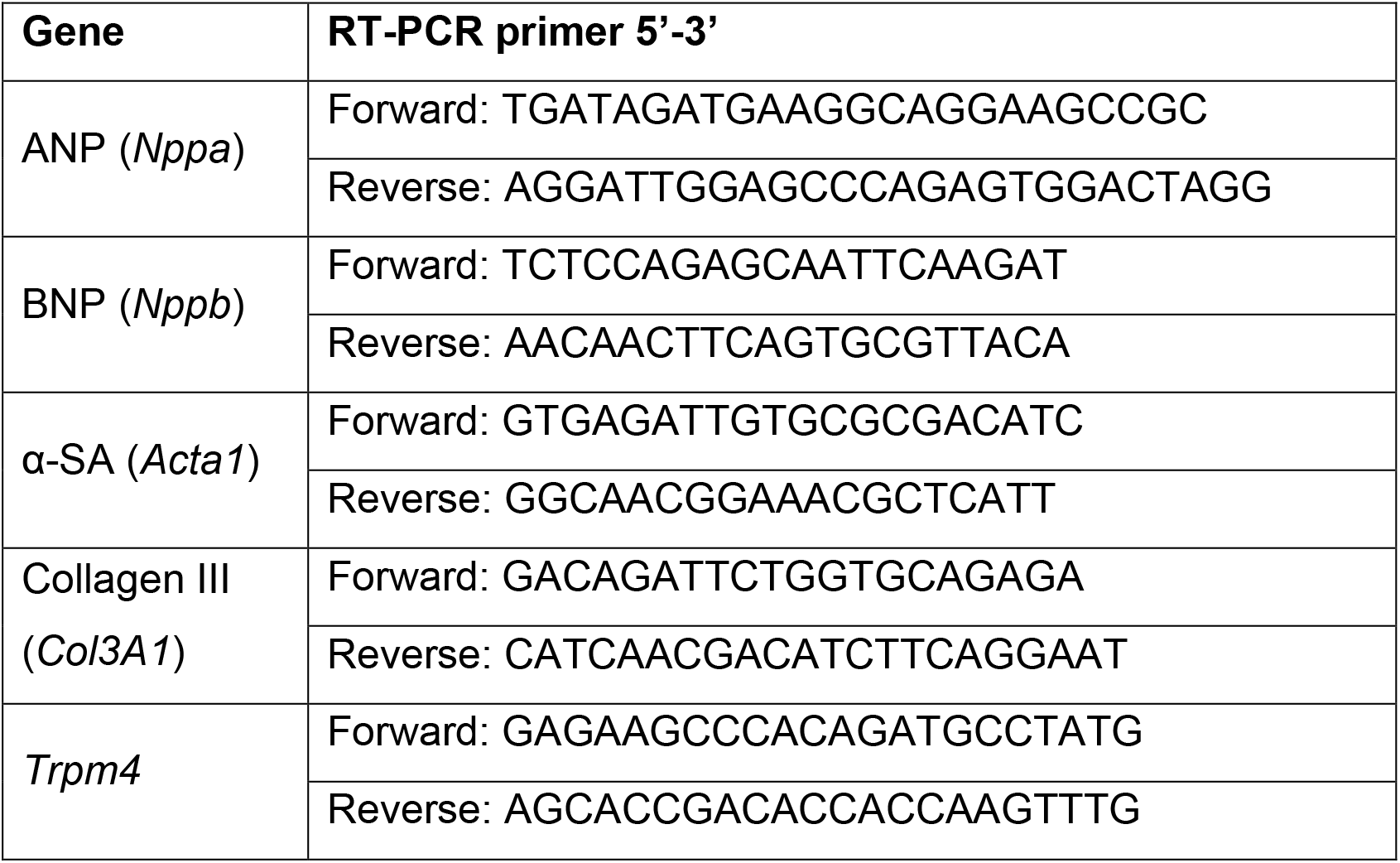
RT-PCR primers

## Notes

### Competing Interest Statement

The authors have declared no competing interest.

